# Hybridization but minimal introgression: ecologically-based divergent selection maintains a steep hybrid zone in parapatric stickleback fish

**DOI:** 10.1101/2025.11.17.688987

**Authors:** Olivia Baebler, Quiterie Haenel, Krista Oke, Andrew P. Hendry, Daniel Berner

**Affiliations:** Department of Environmental Sciences, Zoology & Evolution, University of Basel, Vesalgasse 1, 4051 Basel, Switzerland; Alaska Pacific University, 4101 University Drive, Anchorage, Alaska, AK 99508, USA; Redpath Museum and Department of Biology, McGill University, Montréal, QC, Canada; Department of Environmental Sciences, Zoology & Evolution, University of Basel, Vesalgasse 1, 4051 Basel, Switzerland.

## Abstract

Steep hybrid zones provide key insights into the mechanisms of speciation by reflecting incomplete reproductive isolation between diverging populations. However, the specific reproductive barriers preventing the fusion of such populations generally remain unclear, particularly the role of ecologically-based divergent selection. To address the latter, we here investigate a steep hybrid zone between lake and stream ecotypes of threespine stickleback fish inhabiting contiguous habitats within a single watershed. Given the spatial proximity of these habitats and the system’s postglacial age, historical allopatry is unlikely to have contributed to the evolution of reproductive isolation. Using individual whole-genome sequencing from clinal sampling sites, we find that hybridization occurs within a narrow zone – just a few hundred meters long – at the transition between lake and stream habitat. Individuals in this contact zone exhibit strongly bimodal genome-wide ancestry, with a rapid shift toward the stream ecotype’s genomic background, consistent with strong selection against lake-derived alleles in the stream habitat. Individual-based simulations tailored to this system demonstrate that divergent ecological selection alone can maintain the observed sharp cline, and illustrate the sustained antagonism between gene flow and selection near the habitat transition. Our findings underscore the power of ecological divergence to generate and maintain reproductive isolation, even in the absence of historical separation, and motivate further empirical work on the ecological underpinnings of steep hybrid zones.

## Introduction

Hybrid zones are geographic regions where genetically diverged groups of organisms meet and produce offspring of mixed ancestry (Barton & Hewitt 1985; Gompert et al. 2017; Stankowski et al. 2021). Most hybrid zones are relatively persistent through time and spatially narrow relative to the dispersal capacity of the organisms involved, hence they likely reflect reproductive isolation between the groups in contact. Because the evolution of reproductive isolation is central to the process of speciation, hybrid zones serve as valuable natural laboratories for studying the origin and maintenance of species (Barton & Hewitt 1985; Jiggins & Mallett 2000; Gompert et al. 2017; Moran et al. 2021).

A central challenge in hybrid zone research lies in disentangling the multiple, potentially interacting components of reproductive isolation. In particular, it is often difficult to assess to what extent hybrid zones are maintained by ecologically-based divergent selection (i.e., extrinsic reproductive isolation) versus reproductive barriers not directly linked to ecology that evolved during historical periods of physical isolation (Bierne et al. 2013; Harrison & Larson 2016; Gompert et al. 2017; Moran et al. 2021; Stankowski et al. 2021). The latter can include intrinsic postzygotic isolation, arising from the independent accumulation of incompatible genetic variants, or behavioral isolation, such as divergence in mating preferences.

In many taxa, steep and persistent hybrid zones involve lineages that diverged over hundreds of thousands to millions of generations in physical isolation before coming into secondary contact (e.g., Harrison 1986; Szymura & Barton 1991; Bell 1996; Teeter et a. 2008; Smith et al. 2013; Dufresnes et al. 2021; Natola et al. 2022; Wang et al. 2022; Kalaentzis et al. 2023; Ebdon et al. 2025; Semenov et al. 2025; but see Stankowski 2013; Westram et al. 2018). While ecological selection may still contribute to reproductive isolation in such cases, the strength and nature of this contribution are often difficult to assess due to confounding with other reproductive barriers (Endler 1977; Barton & Hewitt 1985; Kruuk et al. 1999; Bierne et al. 2013; Moran et al. 2021; Stankowski et al. 2021). A direct opportunity to examine the role of ecological selection, however, arises in hybrid zones involving recently diverged populations where substantial evolution in isolation can be ruled out.

With this opportunity in mind, we here focus on a hybrid zone between parapatric lake and stream ecotypes of threespine stickleback fish (*Gasterosteus aculeatus*). Across the species’ Holarctic range, neighboring lake and stream stickleback frequently show marked phenotypic divergence, reflecting adaptation to limnetic and benthic ecological niches, and this ecological divergence is often accompanied by variable degrees of reproductive isolation and associated genetic differentiation (Reimchen et al. 1985; Lavin & McPhail 1993; Reusch et al. 2001; Hendry & Taylor 2004; Berner et al. 2008, 2009; Deagle et al. 2012; Ravinet et al. 2013; Feulner et al. 2015; Roesti et al. 2015; Moser et al. 2016; Stuart et al. 2017). However, despite their occurrence in close physical proximity and the associated potential for gene flow, little is known about the extent of hybridization between lake and stream stickleback ecotypes at the genomic level.

To address this gap, we recently initiated a genomic investigation (Haenel et al. 2021) of lake and stream stickleback occurring in the Misty Lake watershed on Vancouver Island, Canada (Fig. 1) (Lavin & McPHail 1993; Hendry et al. 2002; Moore et al. 2007; Hanson et al. 2016). This ecotype pair has been revealed by previous ecological, biogeographic and phylogenetic evidence to be postglacial in origin (<12,000 generations) and to have diverged in parapatry (i.e., primary intergradation) (Hendry & Taylor 2004; Stuart et al. 2017; Haenel et al., 2021). Experimental crosses between Misty lake and stream fish reveal no intrinsic postzygotic incompatibilities (Lavin & McPhail 1993; Hendry et al. 2002; Berner et al. 2011; Poore et al. 2023) – a general result for postglacial stickleback populations (Hendry et al. 2009) – and mating trials indicate negligible genetically-based sexual isolation between the ecotypes (Raeymaekers et al. 2010; Räsänen et al. 2012). Nevertheless, our clinal genomic work based on pooled whole-genome sequencing uncovered a steep genome-wide transition in allele frequencies over a short (c. 200 m) spatial scale, coinciding with the stream section immediately adjacent to the transition from lake and swamp to stream habitat (Haenel et al. 2021). This pattern strongly suggests that divergent selection plays a dominant role in maintaining reproductive isolation. However, the pooled nature of the genomic data limited inferences about the extent of individual-level hybridization and introgression.

**Fig. 1.**
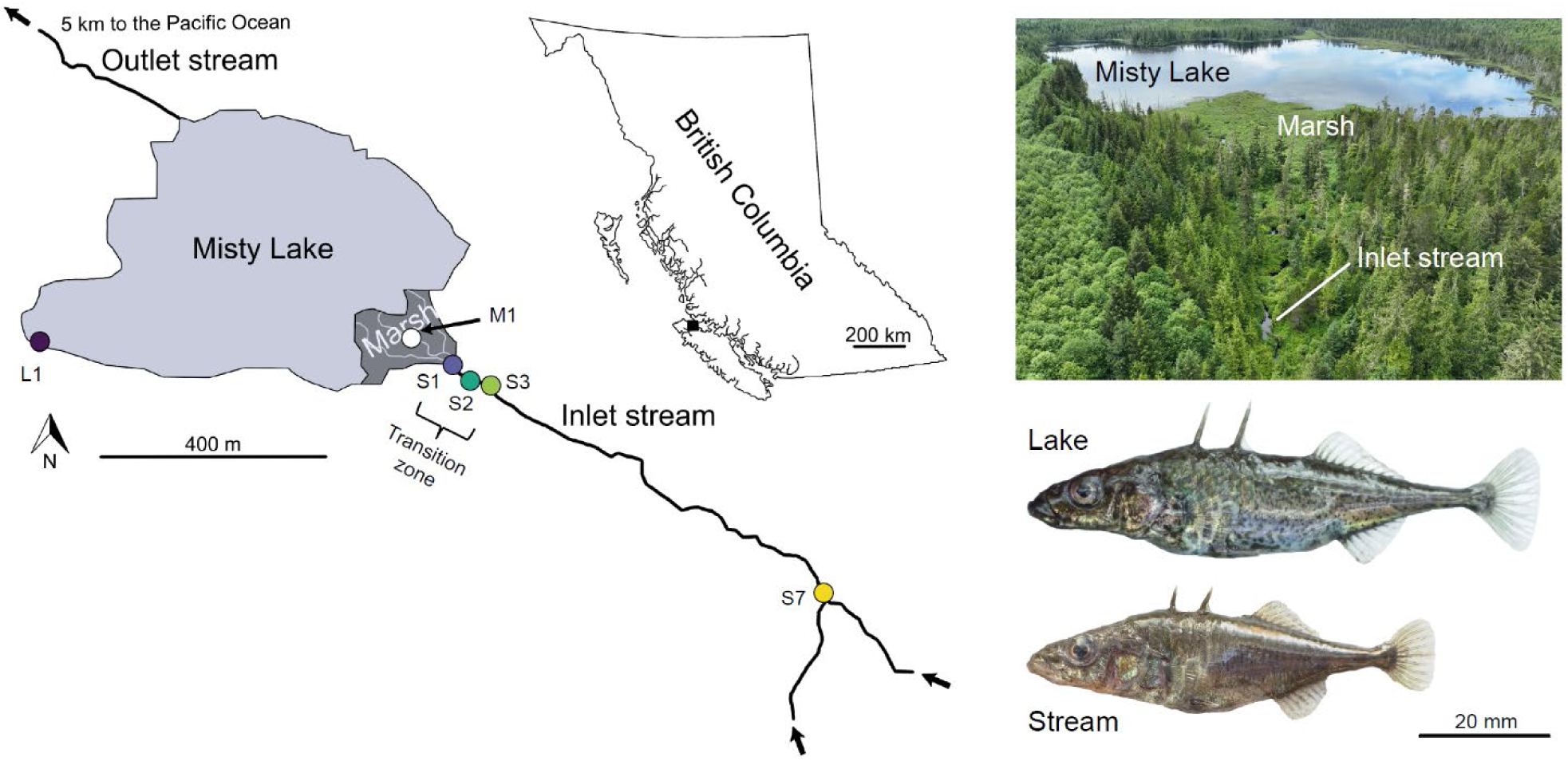
Overview of the Misty Lake and inlet stream system. The dots in the map indicate the lake (L1), marsh (M1) and stream sites (S1-S3, S7) where the experimental individuals were collected (site designation follows Haenel et al. 2021). The insert map shows the location of the Misty watershed on Vancouver Island, British Columbia, Canada. The aerial photograph shows the lake, the marsh, and the lowest reaches of the inlet stream where the genetic transition zone (sites S1-S3) is located. The stickleback specimens are representative males in breeding dress from the pure lake and stream ecotypes. Maps adapted from Haenel et al. 2021, drone image from Andrew Hendry, stickleback specimens from Daniel Berner.

Here, we build on this earlier work by analyzing whole-genome sequences from individually sampled stickleback collected across the previously identified transition zone. Our individual-based genomic analysis reveals clear evidence of hybridization between the lake and stream ecotypes, but at the same time strong genome-wide restrictions to introgression. We support these empirical findings with individual-based simulations tailored to the ecological and spatial characteristics of the Misty lake-stream system. Together, our results provide compelling evidence for the efficacy of ecological selection in maintaining reproductive isolation in a young hybrid zone.

## Materials and Methods

### Study individuals and DNA sequencing

The clinal study by Haenel et al. (2021) using pooled sequencing of stickleback collected during the 2016 breeding season from Misty Lake and its inlet stream identified a pronounced shift in genome-wide allele frequencies occurring over a distance of approximately 200 meters in the lowest reaches of the stream (sites S1, S2, and S3 in Fig. 1; all site names correspond to those in Haenel et al. 2021). Our present investigation of hybridization and admixture is focused on this “transition zone” between the ecotypes and reuses DNA samples from 20 individuals from each of the three transition zone sites (Table S1; see Haenel et al. 2021 for detailed geographic information, and for specimen sampling and DNA extraction methods). To represent the “pure” lake and stream ecotypes with minimal potential hybrid influence, we also included DNA samples from ten individuals collected from locations distant from the transition zone: one site in the lake (site L1) and another in the upper reaches of the stream (site S7) (Fig. 1).

The previous clinal study additionally suggested that the marsh habitat situated between the lake and the inlet stream harbors stickleback genetically slightly differentiated from the true lake population (see also Hanson et al. 2016), with this distinctiveness occasionally disappearing due to pulses of intense dispersal from the lake (Haenel et al. 2021). To verify these observations, we additionally included samples from ten individuals collected at two different time points from the marsh site (M1, Fig. 1). Our main analytical focus, however, is on the transition zone in the stream, as this is where the major genetic shift between the ecotypes occurs. In total, the present work comprises 100 individuals characterized in Table S1.

All individuals were initially subjected to whole-genome sequencing to 101 or 151 bp paired-end reads without enrichment, using three S4 lanes on the NovaSeq 6000 platform of the Genomics Facility Basel, D-BSSE, ETH Zürich. For 12 individuals, this sequencing effort yielded a read depth substantially below our target of 20x. DNA from the latter individuals was therefore enriched through ten amplification cycles and sequenced to 250 bp paired-end reads on an SP flow cell (details provided in Table S1). Final median genome-wide read depth ranged from 10-45x across individuals, with a grand median of 29x.

### Marker ascertainment and genotyping

Raw sequence reads were aligned to the fifth-generation assembly of the stickleback genome (447 Mb; Nath et al. 2021) using NovoAlign (v3.03.00, http://www.novocraft.com/prod ucts/novoalign; parameters settings provided in Code S1). Alignments were converted to BAM format using the R package *Rsamtools* (v2.10.0; Morgan et al. 2022), and base counts were generated at all genome-wide base positions with the *pileup* function (settings provided in Code S1). Using the alignments, we determined the sex of each individual by dividing the number of reads mapping to the X chromosome (Chr19) by the number of reads mapping to an autosome of similar length (Chr20). This ratio ranged between 0.99-1.04 in the homogametic females and between 0.57-0.60 in the heterogametic males.

For single-nucleotide polymorphism (SNP) marker ascertainment, we reused pooled sequencing data from Haenel et al. (2021) from the lake site L1 and the stream site S7, also aligned as described above. These datasets are derived from large samples (*n* = 62 individuals for L1; *n* = 50 for S7) sequenced to high read depth (103x and 107x), thus enabling highly robust SNP detection. Two different SNP panels were ascertained: first, to obtain “ecotype-distinctive SNPs”, we searched for positions exhibiting an absolute allele frequency difference (*AFD*; Berner 2019) of at least 0.85 between the lake and stream sample. This *AFD* threshold, capturing positions where lake and stream fish were highly differentiated (albeit generally not fully fixed for alternative alleles), was chosen to balance physical marker resolution and ecotype specificity. As a robustness check, key analyses were repeated with a more stringent *AFD* threshold of 0.95. Obtaining very similar results throughout, we report only results based on the 0.85 threshold. For the second SNP panel, no *AFD* filter was applied, thus including any level of lake-stream differentiation. These “random SNPs”, however, were still required to exhibit a global minor allele frequency of at least 0.05 across the two pools to minimize sequencing artifacts. At both the ecotype-distinctive and random SNPs, nucleotide counts for both alleles were performed for all our focal individuals. After excluding SNPs located outside the 20 autosomes (i.e., on the sex chromosomes, unassigned scaffolds, or mitogenome), the ecotype-distinctive and random SNP panels comprised 14,929 and 3.84 million SNPs.

### Analysis of overall genetic structure

To gain first insights into the potential hybrid nature of stickleback in the transition zone, we characterized the genetic structure among our study individuals based on a phylogram and an analysis of global ancestry, both using the ecotype-distinctive SNPs. For the phylogram, we derived a haploid FASTA file from the marker panel by concatenating for each individual the most common nucleotide at each SNP (Berner 2021). Using the R packages *ape* (v5; Paradis & Schliep 2018) and *phangorn* (v2.5.5; Schliep 2011), we then generated a maximum likelihood genealogy based on the most likely substitution model of sequence evolution (TVMe + G), and visualized this genealogy as unrooted phylogram.

Global (i.e., overall genome-wide) ancestry was examined by using STRUCTURE (v2.3.3; Pritchard et al. 2000). Input data were derived from the SNP panel by first performing diploid genotype calls for each individual and SNP. For this, we required a read depth of at least 10x but below 2.2x the genome-wide median read depth, and treated variable positions as heterozygous when a binomial test yielded a *p*-value below 0.01 (a threshold determined by preliminary exploration to optimize the detection of true heterozygotes). SNPs with more than 25% missing data across individuals were excluded, resulting in 14,849 markers. Using the same conventions, we additionally generated three replicate input files from the random SNP panel, each independently thinned at random to around 15,000 markers to approximate the marker number of the input data derived from the ecotype-distinctive SNPs. For the ecotype-distinctive data set, STRUCTURE was run with the number of populations *K* = 1-4 in 20 replicates each, with a burn-in period of 100,000 and 200,000 MCMC repeats. The most likely number of populations across our individuals was inferred from the mean likelihood of the data given *K* across runs, and the Δ*K* statistic (Evanno et al. 2005; Earl & vonHoldt 2012). Obtaining unambiguous indication of two populations in this way (see below), the analyses based on the random SNPs were performed with *K* = 2 only.

### Hybridity and heterozygosity

Next, we examined global hybridity and between-ecotype heterozygosity for each individual – metrics capturing information about the depth of admixture in hybrid individuals (Fitzpatrick 2012; Gompert et al. 2017). Using the diploid genotype data derived from the ecotype-distinctive SNPs, an individual’s hybridity was expressed as the proportion of “stream alleles” (i.e., the alleles in high frequency in the stream ecotype) among the total alleles across all genome-wide SNPs. Hybridity near zero thus indicated pure lake ecotype individuals whereas hybridity near one identified pure stream ecotypes. Heterozygosity was quantified as the proportion of heterozygous markers among all genome-wide SNPs for a given individual, a metric for which values near the maximum of one were expected for F1 hybrids between the pure ecotypes. For visual analysis, we plotted heterozygosity against hybridity for all sample sites, and additionally for all transition sites combined. Since the markers in our ecotype-distinctive SNP panel were not perfectly ecotype-diagnostic (i.e., not fully fixed for alternative allele between the L1 and S7 pools), we additionally calculated hybridity and heterozygosity for ten synthetic F1 hybrids, serving as an expectation for that hybrid class. These individuals were generated in two alternative ways: first, we used the Haenel et al. (2021) pooled sequencing data to combine at each ecotype-distinctive SNP a single allele drawn at random from the L1 nucleotide pool according to the observed frequencies of the two SNP alleles, and a single allele drawn similarly from the S7 pool. In a second approach, we formed ten unique L1-S7 individual pairs based on our individual sequencing data and combined at each ecotype-distinctive SNP a single nucleotide drawn from each individual. Because synthetic F1 hybrids derived from pooled versus individual-level sequencing were visually indistinguishable in their hybridity and heterozygosity, we just present results from the latter approach.

### Local ancestry inference

While the above analyses provided global (i.e., genome-wide) insights into hybridization, we additionally took advantage of our whole-genome sequencing resolution to explore potential consequences of hybridization locally along chromosomes within each individual. For this, we used Ancestry_HMM (v1.0.2; Corbett-Detig et al. 2017), a program estimating ancestry probabilities at each genomic position for each individual in a set of samples, plus the age of hybridization based on the simplifying assumption of a single punctuated hybridization event. To generate input data, we considered only markers without any missing data across all study individuals and exhibiting a minor allele frequency of 0.25 or greater in the combined L1-S7 sequence pool, and thinned these SNPs to ≥ 2 kb physical spacing to reduce ancestral linkage disequilibrium (LD), resulting in 153,065 genome-wide SNPs. (Strong within-population LD typically decays over a few kilobases in this species, e.g., Roesti et al. 2015). Because LD-thinning may influence the age estimation of hybridization, we additionally considered input data based on ≥ 5 kb marker spacing (70,310 SNPs). However, due to unavoidable deviation of our empirical system from model assumptions (e.g., a single hybridization pulse, Corbett-Detig et al. 2017), we interpret the estimated hybridization ages as qualitative indications. Ancestral SNP allele counts required by the software were determined for each marker by treating L1 and S7 as reference populations and estimating allele frequencies from their sequence pools. Average global admixture proportions, also required as input information, were available from our STRUCTURE analysis (as performed with the random SNPs). Genetic map distances between adjacent markers were calculated based on their physical distance and assuming a uniform crossover rate of 3.11 cM/Mb (Roesti et al. 2013). The simplifying assumption of a uniform recombination rate across the genome is not expected to materially influence local ancestry inference, nor the age estimation of hybridization (Corbett-Detig et al. 2017; R. Corbett-Detig, personal communication). Within-individual variation in ancestry was visualized by local ancestry probabilities (i.e., homozygous for lake or stream ecotype ancestry, or heterozygous) along the first three chromosomes for selected individuals from all sites.

Focusing on the transition zone only, we next asked if specific genome regions were particularly enriched for stream ancestry, potentially revealing genome regions under selection for stream alleles. Since the transition zone proved to harbor a mix of direct lake migrants (or weakly admixed lake ecotype-like individuals) and more strongly admixed, advanced-generation hybrid individuals (see Results), we excluded the former, as only in the latter the opportunity for the selection of localized genomic regions was given. The transition zone individuals considered as admixed and retained for this analysis were required to exhibit a hybridity greater than 0.2 (*n* = 35, Table S1). Across these individuals, we averaged the estimated probability of homozygous stream ancestry for each marker, and visualized the resulting values along the first three chromosomes.

To examine a potential footprint of selection in the transition zone more formally, we identified all markers with an exceptionally high (>= 0.8) average probability for homozygous stream ancestry, and assessed if this small subset of markers (*n* = 736 SNPs located on 15 chromosomes) also displayed elevated differentiation (*AFD*) between the L1 and S7 sample based on the pooled sequence data. Such an association would suggest that genome regions harboring alleles locally favored in the transition zone tend to coincide with those under long-term divergent selection between the pure ecotypes. As a control, we repeated this procedure based on markers exhibiting an exceptionally high (>=0.5) average probability for homozygous *lake* ancestry, expecting no elevated L1-S7 pooled sequencing differentiation for this SNP subset (636 markers on 14 chromosomes). The benchmarks for these analyses were generated by bootstrap resampling the full set of markers used for ancestry analysis 10,000 times to the same size as the high stream and high lake ancestry subsets, and for each iteration’s subsets recalculating average ecotype differentiation from the pooled sequencing data. To verify the robustness of the results obtained, we modified this analysis of signatures of selection by using 10 kb sliding windows instead of individual SNPs as data points, the median instead of the mean as location statistic, and more or less stringent ancestry probability thresholds for marker subset selection. None of these modifications led to qualitatively different results (details not presented).

### Influence of recombination rate variation on introgression

Divergent selection between lake and stream stickleback is polygenic (Roesti et al. 2012, 2015; Rennison et al. 2019; Haenel et al. 2021; Laurentino et al. 2021; Poore et al. 2023), and recombination rates in stickleback are relatively consistently and strongly elevated in the chromosome peripheries compared to the chromosome centers (Roesti et al. 2012, 2013).

Combined, these conditions may allow for heterogeneous introgression between habitats across the genome (Berner & Roesti 2017; Haenel et al. 2018; Veller et al. 2023). Specifically, under divergence with gene flow and associated admixture, chromosome segments containing locally advantageous alleles may be especially long in chromosome centers where recombination rates are relatively low. In these regions, divergent selection may be particularly effective due to the strong physical linkage of alleles, potentially leading to an enrichment of locally adaptive ancestry in chromosome centers (Roesti et al. 2012; Berner & Roesti 2017; Veller et al. 2023).

We explored such potential influence of broad-scale heterogeneity in recombination rate on introgression both for the transition zone, and for the pure populations. For the former, we averaged individual probabilities of pure stream ancestry at each SNP across all individuals from the transition zone, again considering only the 35 individuals substantially admixed (Table S1). We then calculated mean stream ancestry across all SNP-specific averages separately for the periphery of each chromosome, and for their centers. Following Haenel et al. (2021), we considered the 5 Mb from either chromosome tip as its periphery, and the rest as its center.

Using the 20 autosomes as replicate data points, we then explored differences in stream ancestry probability between chromosome centers and peripheries. For this, we expressed the 95% compatibility intervals for the chromosome region-specific median ancestry probabilities by the central 95 percentiles of the bootstrap distributions based on 10,000 bootstrap re-samples of the chromosomes. Because our L1 and S7 samples included only ten individuals, we used differentiation (*AFD*) between the sequence pools along chromosomes instead of local ancestry to examine heterogeneous introgression between the pure ecotypes. Apart from this modification, we followed the same analytical protocol as for the transitions zone.

Introgression probability along chromosomes may not only be influenced by heterogeneity in recombination rate, but also by heterogeneity in gene density. Gene-rich chromosome regions, likely exhibiting an elevated density of selection targets, may be selected for locally favorable ancestry more effectively and hence show relatively reduced introgression.

We therefore considered this potential confounding factor by calculating gene density (number of genes per Mb) for the peripheries and centers of all autosomes, based on the gene annotation of the stickleback genome (Nath et al. 2021). This revealed no relevant difference in gene density between stickleback chromosome centers and peripheries (Fig. S1), so that we did not further consider this factor in the above analyses.

### Individual-based simulations

To improve our understanding of the key determinants of hybridization and admixture at the lake-stream transition, we complemented our empirical analyses by individual-based simulations performed with a modification of the stepping-stone model developed for the Misty lake and inlet stream system in Haenel et al. (2021). While the original model served to examine changes in pooled allele frequencies along a linear array of adjacent demes, the present implementation aimed to identify combinations of key population genetic parameter values able to generate genetic signatures in the transition zone similar to the ones observed empirically. The model comprised a total of nine demes, all of which harboring a total of 200 diploid individuals except for the first “lake” deme. The latter was, following empirical evidence (Fisheries and Ocean Canada 2018), modeled *f* times larger than each of the other (“stream”) demes. The first stream deme, directly adjacent to the lake deme, was our focal transition zone deme. Based on ample evidence of polygenic divergent selection on stickleback in lake and stream habitats, we modeled 100 unlinked biallelic loci under selection. At each locus, one allele was favored in the lake deme but deleterious in all stream demes, while the opposite was true for the alternative allele. In the beginning of all simulations, the two alleles were sampled at random with the same probability of 0.5 across all individuals and demes to avoid stochastic allele loss during the early generations. The model scenario thus mimicked primary adaptive divergence from standing genetic variation typical of stickleback fish (Jones et al. 2012; Terekhanova et al. 2014; Lescak et al. 2015; Roesti et al. 2015; Bassham et al. 2018; Haenel et al. 2019; Galloway et al. 2020).

An individual’s performance within a given deme was determined by its genotype across all loci, assuming a relatively weak per-allele (haploid) selection coefficient of 0.005 (reduction in performance from the local optimum of 1). Each generation involved a migration phase during which each deme sent a fraction *m* of its individuals drawn at random into each adjacent deme. This was followed by a phase of selection and reproduction during which a deme’s offspring cohort was generated by drawing two individuals from the parental cohort at random, but weighted by their performance, to produce a single offspring until initial deme size was re-established. Individuals were here treated as hermaphrodites and allowed to mate multiple times. All simulations were run over 1000 generations, although the model already reached migration-selection equilibrium after around 500 generations (Fig. S2).

To keep the simulation effort within reasonable limits, we initiated our analysis by a preliminary series of runs in which we haphazardly varied model parameter values and looked for patterns of hybridity and heterozygosity in the transition zone deme qualitatively resembling the ones observed empirically for the real transition zone. This preliminary screen (details not presented) identified two parameters as particularly relevant: the scaling factor of the size of the lake deme relative to the stream demes, as well as the migration proportion (i.e., the parameters *f* and *m* described above). We thus explored these two parameters in detail by using Approximate Bayesian Computation (ABC). For this, we performed 300 runs with our simulation model, each time using a unique combination of parameter values drawn at random from uniform distributions bounded between 2-50 for *f* and 0.01-0.2 for *m*. With the largest parameter values considered, the model thus comprised 11,600 total individuals, and 20% of a deme’s individuals migrated into each adjacent deme (i.e., 40% emigration for the non-terminal demes, 20% for the terminal ones).

At generation 1000, we characterized the transition zone deme after migration but before reproduction (i.e., the parental cohort, corresponding to our empirical samples). We here took a subsample of 20 individuals to match our empirical sample size, calculated median hybridity and heterozygosity for the five individuals with the highest and lowest values for each variable plus for the subsample as a whole, and saved the resulting six statistics from each run along with the associated values for *f* and *m*. Next, we calculated the same hybridity and heterozygosity statistics for the real sample from site S1, serving as target values for ABC estimation. Finally, we ran 100 replicate ABC estimation runs using the R package *abc* (v2.2.2; Csillery et al. 2012) with the neural network algorithm and a tolerance of 0.1 to identify values of *f* and *m* for which the simulated statistics best approximated the empirical targets (“estimation optima”). In addition, we ran our simulation model with the most plausible *f* and *m* values identified in this way in ten replicates to quantify patterns of hybridity and heterozygosity at generation 1000, both after migration but before reproduction (parental cohort), and after reproduction but before migration (offspring cohort). This allowed us to appreciate changes in genetic composition occurring within a generation. Finally, to explore the influence of the *f* and *m* parameters in isolation, we ran simulation series in which we varied just one of these parameters across five levels (*f* = 3, 6, 12, 25, 50; *m* = 0.01, 0.02, 0.05, 0.1, 0.2) while keeping the other parameter at ABC optimum.

Unless specified otherwise, all analyses, graphing and simulations were implemented with the R language (R Core Team 2024).

## Results and discussion

### The transition zone is influenced by both dispersal and hybridization

The phylogram based on the ecotype-distinctive SNPs revealed that the vast majority of the 60 total individuals collected from the transition zone (Fig. 1) grouped together either with the pure lake individuals (most of the S1 sample), or with individuals from the upper stream (the majority of the S2 and S3 samples), thus indicating strong bimodality in genomic composition within this stream section (Fig. 2a). A few individuals from the transition zone, however, branched between these two deeply separated groups, suggesting the presence of hybrids. Global population ancestry analysis complemented this tree-based insights by confirming the presence of two distinct genetic populations (Fig. S3) in the Misty lake-inlet stream system (consistent with the microsatellite-based result in Moore et al. 2007), and of a small number of admixed individuals with intermediate ancestry (Fig. 2b). The latter proved mostly restricted to the lowest stream site S1; most transition zone individuals either exhibited an ancestry composition similar to the lake fish, or approached the ancestry composition seen at the upper stream site. Using for global ancestry inference the random SNPs instead of the ecotype-distinctive ones produced very similar results supporting the same conclusions (Fig. S4).

**Fig. 2.**
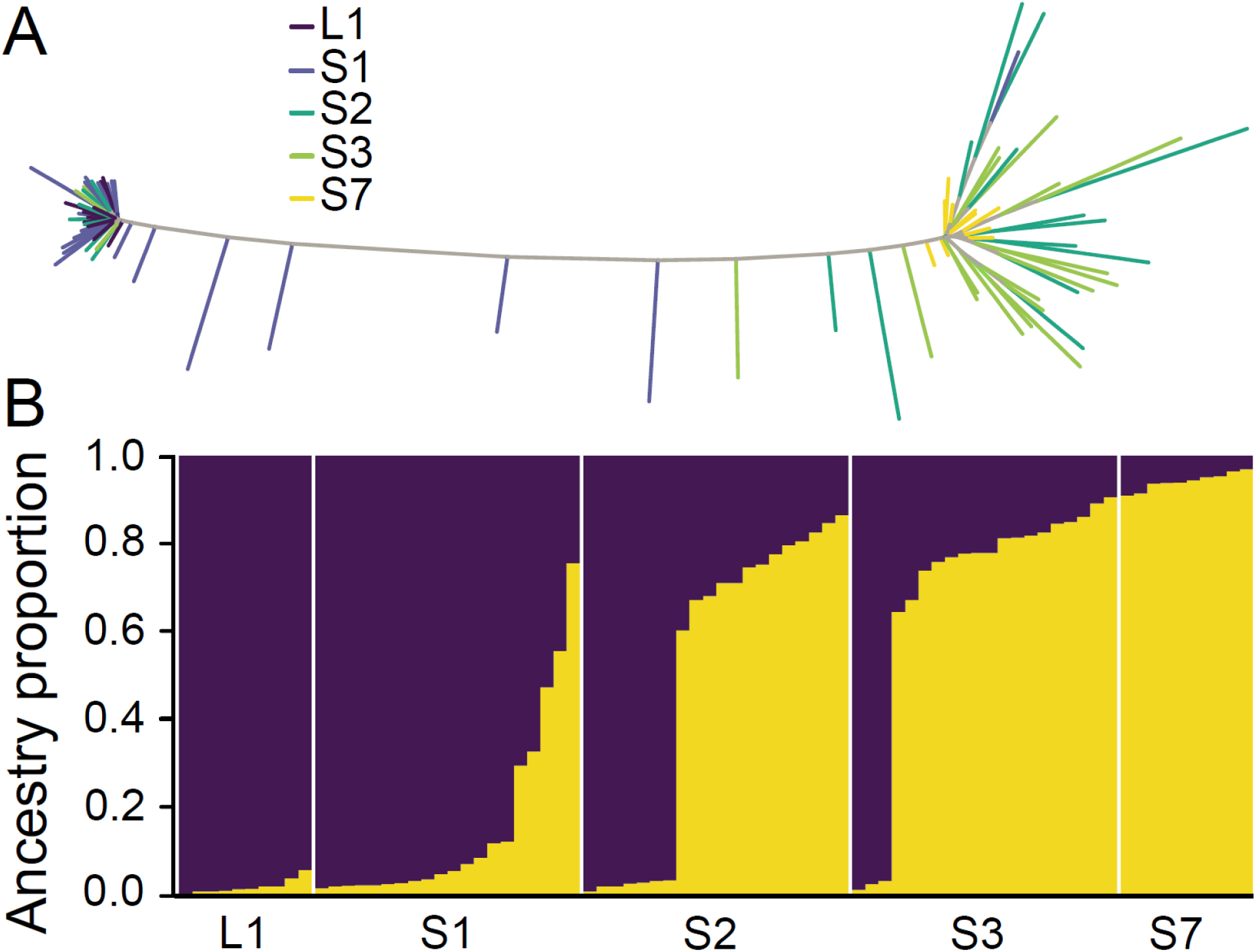
Population structure among stickleback from Misty Lake (L1), the upper inlet stream (S7), and the transition zone (S1-S3), based on the ecotype-distinctive marker panel. In A), individuals are visualized in an unrooted phylogram, color-coded by sample site according to Fig. 1. B) Global ancestry, sorted by increasing stream ecotype ancestry proportion within each sample site (individuals are represented by columns).

Previous work on Misty lake and inlet stream stickleback based on pooled sequencing identified a sharp transition in marker allele frequencies in the lowest reaches of the inlet stream (Haenel et al. 2021). This raised the question to what extent this transition reflects hybridization and associated genomic admixture, as opposed to just dispersal causing genetically differentiated, non-admixed ecotypes to co-occur. Our current examination of population genomic structure across the transition zone using individual-level sequence data resolves this question by indicating the presence both processes: on the one hand, we observe in the transition zone a small fraction of individuals genomically intermediate between the lake and the stream ecotype, and a larger fraction of individuals displaying a proportion of stream ancestry approaching – yet still substantially below – the one seen in the pure stream ecotype sample. All these individuals are clearly admixed, hence must derive from hybridization. On the other hand, we find a substantial fraction of individuals genomically very similar to, or indistinguishable from, the lake ecotype. We interpret these individuals as direct dispersers from the lake. This dispersal is highly restricted spatially, as we observe a strong decline in the fraction of such lake dispersers from the site S1 to S3, that is, over a stream section of just around 90 meters (Fig. 1). That lake dispersal may indeed not reach much beyond the transition zone is consistent with the absence of pooled allele frequency changes across the whole stream section from a site located around 80 meters upstream of our site S3 up to the site S7 (Haenel et al. 2021).

### The transition zone harbors a swarm of deeply admixed hybrids

Having demonstrated hybridization in the transition zone, we aimed to examine the depth of genomic admixture resulting from hybridization, and the consequences of hybridization to ancestry at the chromosome scale. To initiate the former, we plotted individual heterozygosity against hybridity based on the ecotype-distinctive SNPs. This polarized the lake and stream ecotypes to opposite hybridity, and both groups exhibited low heterozygosity (Fig. 3, top), as expected for markers strongly but not perfectly ecotype-diagnostic. At all transition zone sites, we observed individuals genetically similar to the lake fish (delimited by a hybridity <0.2), with a sharp decline in their fraction from S1 (75%) to S2 (35%) and S3 (15%) (Fig. 3, middle rows).

**Fig. 3.**
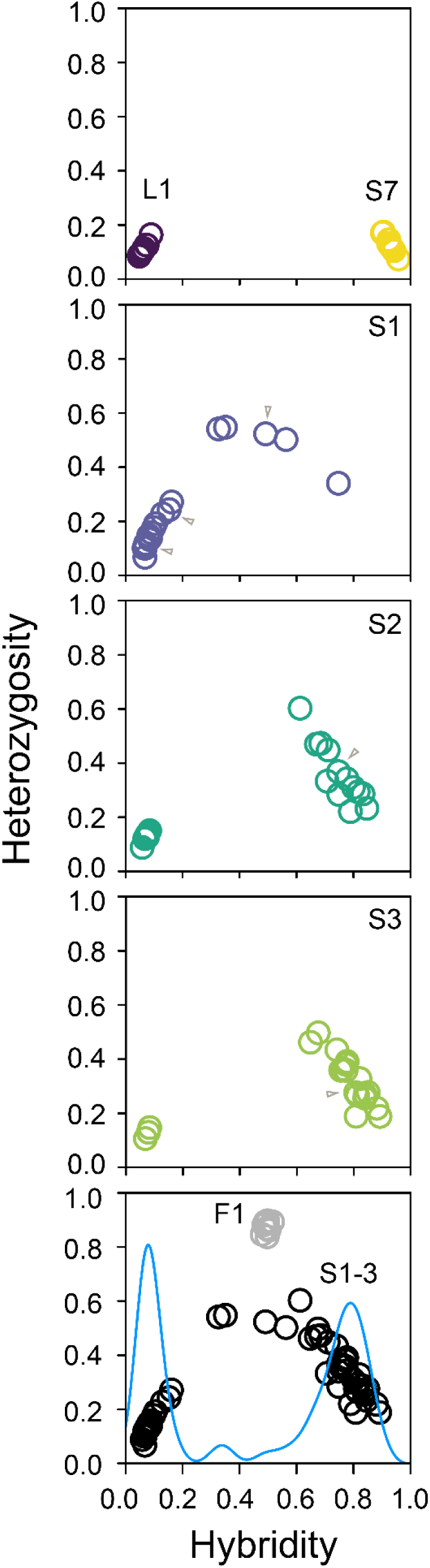
Individual hybridity and heterozygosity at the sample sites, based on the ecotype-distinctive SNPs. Hybridity represents the proportion of stream alleles across all markers, while heterozygosity expresses the proportion of SNPs heterozygous for lake and stream alleles. The small gray arrows indicate individuals for which local ancestry along chromosomes is presented in Fig. 4. The bottom panel depicts all individuals from the transition zone together, with hybridity kernel density-smoothed (band width 0.04) to highlight bimodality. The same panel additionally indicates values expected for F1 hybrids between pure lake and stream ecotypes, based on ten synthetic hybrid individuals.

These individuals were potentially male-biased; only a third (eight out of 25) were females, although statistical precision was low (95% CI for female probability: 0.15-0.54). The site S1 harbored a few individuals with intermediate hybridity, whereas the majority of individuals from the sites S2 and S3 tended toward the signature of the stream ecotype. Across the transition zone as a whole, hybridity was thus strongly bimodal, and individuals consistent with F1 hybrids were notably absent (Fig. 3, bottom).

These observations based on allele frequencies at the ecotype-distinctive SNPs suggested relatively deep admixture (i.e., the mixing of the pure ecotypes’ genomes over multiple generations) for most of the transition zone individuals. This was examined further by inferring local ancestry along chromosomes from the random SNPs. As expected, this revealed relatively homogeneous genome-wide ancestry in fish from L1 and S7 (Fig. 4a, top and bottom row). A fraction of individuals from the transition zone, in particular from S1, showed ancestry patterns similar to those from the lake sample (Fig. 4a, second row), clearly identifying them as migrants from the lake. The other fish from the transition zone generally exhibited numerous ancestry shifts along their chromosomes (Fig. 4a, rows 4-6), consistent with a deep admixture history involving extensive recombination between typical lake and stream ecotype chromosomes. Individuals from the transition zone were indeed estimated to originate from admixture pulses between the pure lake and stream ecotypes having occurred between 151 (S2) and 322 (S1) generations ago. Although this result was somewhat contingent on our marker thinning (with 5 kb-thinning, the pulses were estimated between 64 and 140 generations in the past), sustained admixture over numerous generations within the transition zone is unequivocal. Interestingly, a few individuals from S1 with slightly higher hybridity than typical lake migrants (around 0.15) showed distinctive tracts of extended heterozygous ancestry on multiple chromosomes (see chromosome 1 in third row of Fig. 4a, and further examples in Fig. S5), suggesting recent backcrossing of admixed individuals to lake migrants.

**Fig. 4.**
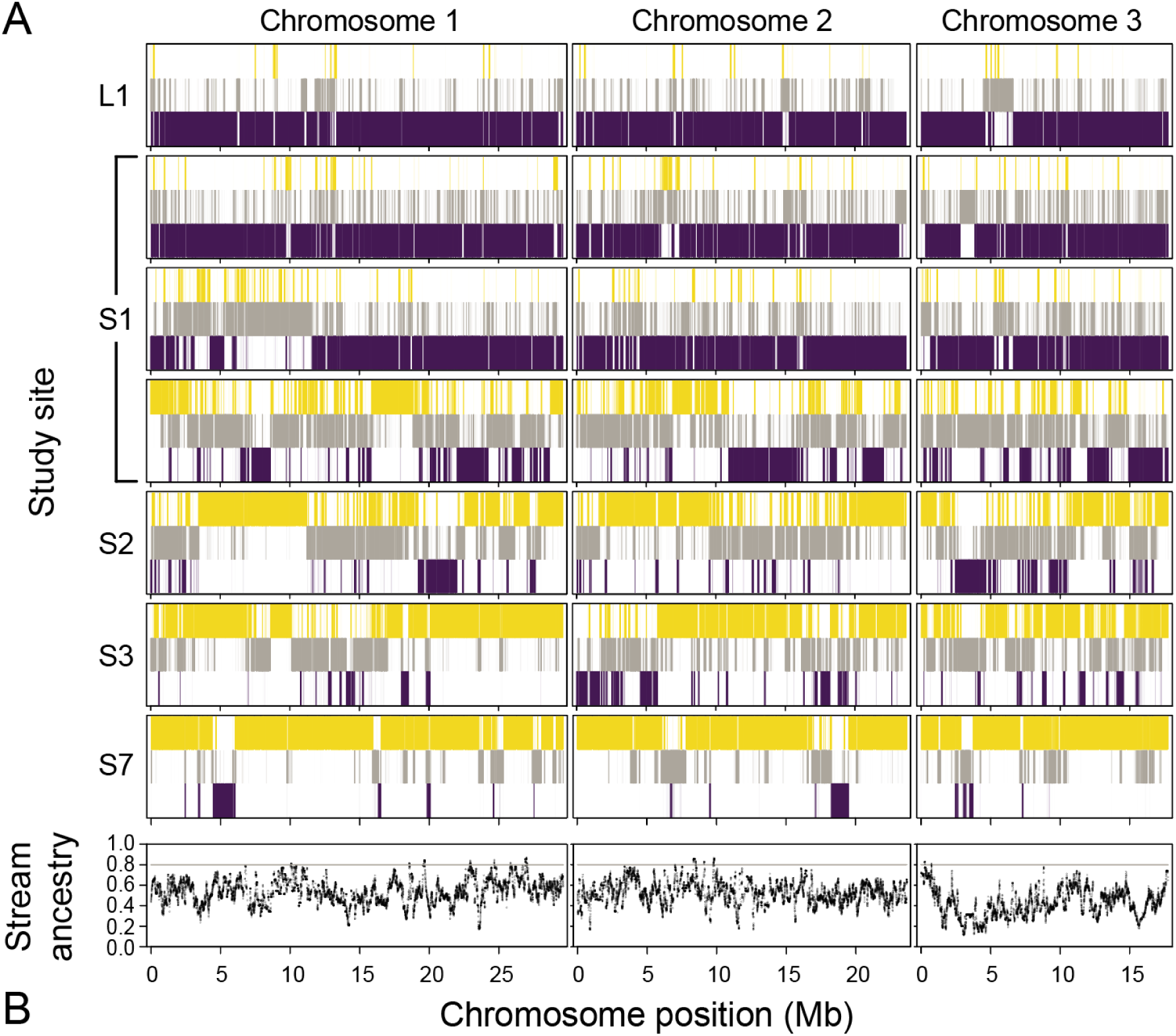
Local ancestry along three representative chromosomes, as inferred from the random SNP panel. In A), probabilities of homozygous lake and stream ecotype ancestry (color coded as in Fig. 2) or heterozygous ancestry (gray) are presented for a pure lake and stream individual (top and bottom row), and for selected individuals (flagged by gray arrows in Fig. 3) from the transition zone (middle rows). For the site S1, a lake disperser, a weakly admixed and a strongly admixed individual are shown. B) Probability of homozygous stream ancestry averaged over the 35 transition zone individuals exhibiting substantial admixture between the ecotypes (hybridity > 0.2, see Fig. 3 bottom panel). The gray horizontal line indicates the threshold chosen to delimit markers with an exceptionally high stream ancestry probability examined for a collective signature of selection.

These analyses of admixture depth make clear that the transition zone is composed of two main classes of individuals: immigrants from the lake, and a swarm of deeply (over many generations) admixed hybrids with predominant stream ancestry. Across the transition zone, hybridization involving pure lake fish appears to occur in immediate proximity to the lake-stream habitat transition only (site S1); the sites S2 and S3, located just a few dozens of meters further upstream, harbor no individuals of intermediate hybridity. Across the latter section of the transition zone, admixed individuals must thus interbreed, but not backcross with the pure lake immigrants, or at least such backcrossing produces no surviving offspring. Because pure stream ecotype individuals are absent across the entire transition zone, F1 hybrids are never produced. Overall, the genomic bimodality seen in stickleback from the transition zone is a clear indication of strong reproductive isolation driven by divergent selection against genomically intermediate individuals (Barton & Hewitt 1985; Jiggins & Mallet 2000; Stankowski et al. 2021). Sexual reproductive barriers are unlikely to contribute to this bimodality, as experimental evidence reveals mating isolation between the Misty lake and stream ecotypes and their hybrids to be weak at best (Raeymaekers et al. 2010; see also Irwin 2020).

### Signature of selection in the transition zone

To obtain evidence of selection in the transition zone, we inspected average local ancestry probabilities along chromosomes across the 35 individuals proving well-admixed (hybridity > 0.2). This revealed high heterogeneity in ancestry probabilities (Fig. 4b), and a clear footprint of selection: markers exhibiting exceptionally high average stream ancestry probability also showed substantially (28%) elevated genetic differentiation between the pure lake and stream ecotype samples shaped by log-term selection (mean *AFD* = 0.468), compared to marker samples of similar size drawn at random (resampling mean *AFD* = 0.367, range 0.333-0.397) (Fig. 5a). Conversely, markers exhibiting a high average *lake* ancestry probability in the transition zone displayed 10% lower average L1-S7 differentiation (*AFD* = 0.332) than random marker samples, consistent with these loci being relatively unimportant to adaptive divergence. Neither ancestry probabilities in the transition zone nor the magnitude of differentiation between the pure ecotypes proved substantially influenced by large-scale heterogeneity recombination rate, as quantified by the central versus peripheral position of markers along chromosomes (Fig. 5b; genomic differentiation between the pure ecotypes is characterized in detail in Haenel et al. 2021).

**Fig. 5.**
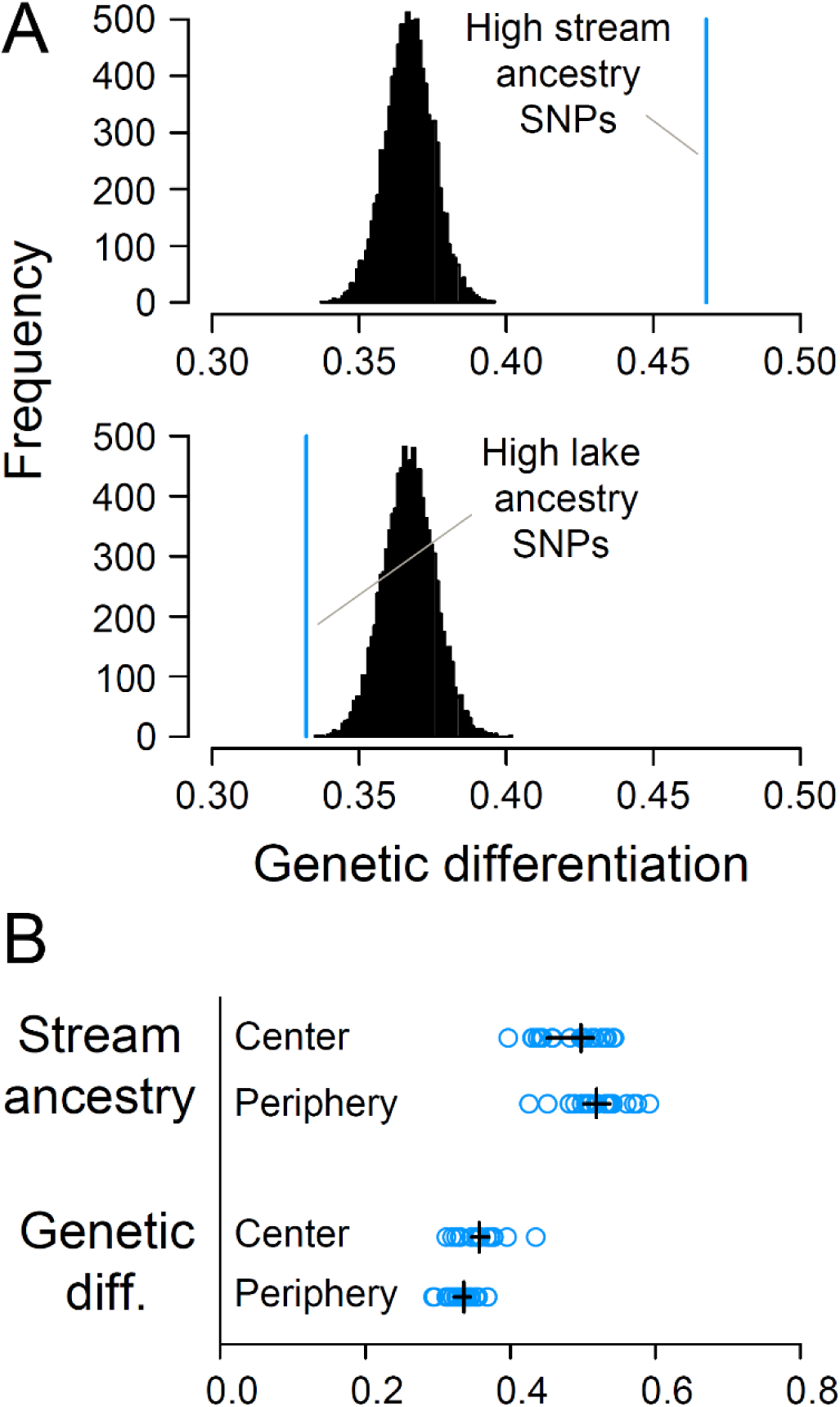
A) Signature of selection in the transition zone. The blue vertical lines indicate the mean magnitude of genetic differentiation (expressed by the absolute allele frequency difference AFD) between the pure lake and stream ecotype pools for markers showing exceptionally high homozygous stream ecotype ancestry (top, *n* = 736 SNPs, see Fig. 4B) and lake ecotype ancestry (bottom, *n* = 636 SNPs) in the transition zone. Only individuals with substantial admixture (hybridity > 0.2) were considered (see Fig. 3 bottom). As baseline for comparison, both graphs include the distribution of the analogous statistic across 10,000 resamples of SNPs with random ancestry. B) Check of an influence of heterogeneity in recombination rate (low in the chromosome centers, high in their peripheries) on the probability of homozygous stream ecotype ancestry in the transition zone (top), and on the magnitude of genetic differentiation (*AFD*) between the pure lake and stream ecotype pools (bottom). The blue circles are chromosome-specific means, the black vertical lines are their medians, and the black horizontal lines are 95% bootstrap compatibility intervals for the latter based on 10,000 chromosome resamples.

The lack of a substantial influence of heterogeneity in recombination rate – known to be strong along the stickleback chromosomes (Roesti et al. 2013) – on ancestry probabilities and lake-stream ecotype differentiation in the Misty system aligns well with theoretical expectations: heterogeneous recombination rate modifies the efficacy of polygenic divergent selection across the genome only when genetic mixing between divergent populations is sufficiently extensive (Berner & Roesti 2017). As implied by the strong bimodality in hybridity across the transition zone, such mixing is largely prevented by divergent selection even in immediate proximity to the lake-stream transition. In other words, the residence time in the stream of locally unfavorable chromosome segments from the lake may be too short for variation in recombination rate along chromosomes to influence introgression probability substantially.

Strong divergent selection, and the associated reproductive isolation between the lake and stream ecotypes, may also offer an explanation for the absence of sexual isolation (mating barriers) between the ecotypes from the Misty system, as found in laboratory experiments (Raeymaekers et al. 2010; Räsänen et al. 2012). Given strong selection against the lake genome in the stream, avoiding hybridization through sexual premating isolation would appear selectively favorable, but the evolution of such a sexual barrier (i.e., reinforcement) requires extensive opportunity for mating and hybridization between the pure ecotypes (Coyne & Orr 2000; Servedio & Noor 2003). As our analyses indicate, these conditions are not met; hybridization occurs too rarely and locally in the Misty system as a whole.

### Simulations support asymmetric ecotype population sizes and massive lake dispersal into the transition zone

Our empirical analyses above indicate a key role of dispersal and divergent selection in the Misty lake-inlet stream system. To explore whether observed patterns of admixture can arise just from a simple antagonism between dispersal and selection, we combined individual-based stepping-stone simulations with approximate Bayesian computation (ABC). This revealed that patterns of hybridity and heterozygosity best approximating our empirical results can be reproduced with a model involving a lake deme much (28x) larger than the stream demes, combined with a migration rate around 0.09 among adjoining demes (Fig. S6). Simulations with these optimal parameter estimates indeed produced 1) strong local adaptation in the terminal demes of the model (Fig. 6A, left); 2) a first stream deme dominated by lake migrants but exhibiting a minor fraction of highly admixed individuals (Fig. 6A, middle); and 3) a strong shift toward predominant stream hybridity in the second stream deme (Fig. 6A, right) (compare to the three upper rows in Fig. 3). Note that the model allowed for dispersal between neighboring demes only, hence stream demes beyond the first one lacked the direct lake migrants observed empirically at the sites S2 and S3. Modelling weaker asymmetry in deme size prevented strong local adaptation in the lake deme, and hence the possibility of dispersal of lake individuals with low hybridity into the first stream deme (Fig. 6B, upper row). Modelling a lower dispersal rate in turn reduced the probability of hybridization and the associated occurrence of admixed individuals in the first stream deme (Fig. 6B, lower row). With an extremely high dispersal rate, however, the first stream deme became overwhelmed by dispersers from the lake, thus impeding local adaptation and hence admixture.

**Fig. 6.**
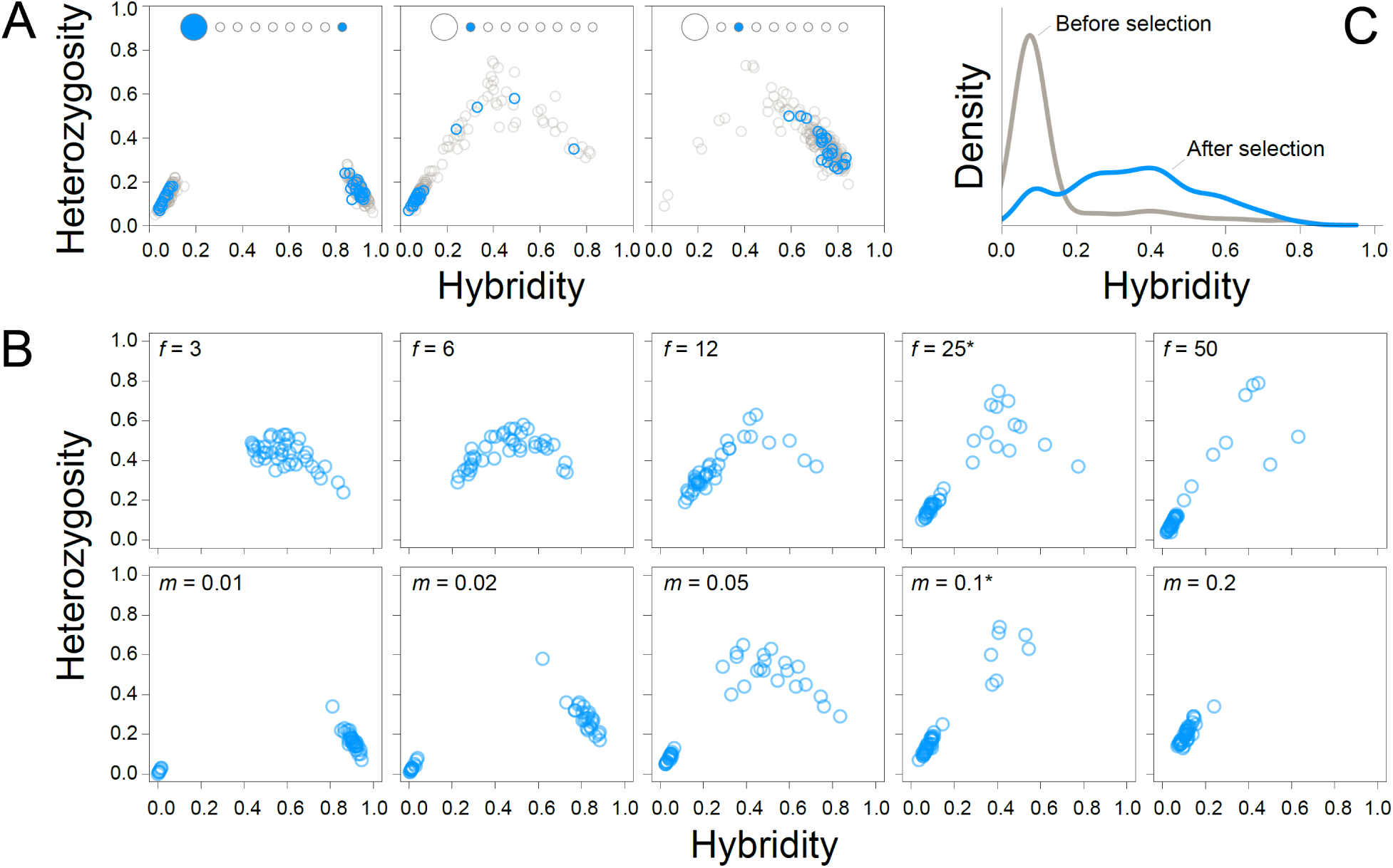
Individual hybridity and heterozygosity in the stepping-stone simulation model at migration-selection equilibrium. In A), results based on ABC parameter optima for the deme size scaling factor (*f* = 28) and the migration rate (*m* = 0.09) are shown for individuals from the lake and the terminal stream deme (left), and for the first and second stream deme (middle and right). The position of the demes in the model is illustrated schematically at the top of each panel. The gray data points represent a pool (*n* = 200) of 20 randomly chosen individuals from each of the ten replicate simulation runs, while the blue data points show a random sample of 20 individuals from a single replicate. B) Influence on the genetic composition in the first stream deme of varying the *f* or *m* parameter in isolation (i.e., keeping the other parameter at ABC optimum). The data points are 40 random individuals from a single simulation run. The parameter values flagged by an asterisk are those closest to their ABC optimum. C) Change in hybridity within a single generation in the first stream deme. The curves reflect kernel density smoothed (band width 0.04) relative density, not absolute individual number (which is much higher before selection due to immigration). Hybridity peaks low before selection because most individuals are fresh immigrants from the lake, but rises strongly due to selection.

Our model with ABC parameter optima further highlighted massive oscillation in the genetic composition of individuals in the stream deme adjacent to the lake during each generation. Due to the large size of the lake deme relative to the stream demes, dispersal caused around 97% of the individuals in the model’s first stream deme to be direct dispersers from the lake with minimal hybridity (Fig. 6C). However, the hybridity of the offspring cohort, that is, after the selection phase, proved greatly shifted upward, thus implying an extremely low local reproductive contribution of the numerically dominant lake migrants across generations. In nature, the latter may occur via massive selective juvenile mortality, which has been suggested for Misty lake-stream stickleback (Hendry et al. 2002) and confirmed for a European lake-stream stickleback system (Moser et al. 2016) by release-recapture experiments. Tracking the genetic composition of stickleback cohorts over time at the site S1 would offer valuable insights into how selection occurs across life stages.

Our simulations, producing patterns of admixture resembling the empirical ones, indicate extensive dispersal from a large lake population into the lowest stream section. Despite this strong and numerically asymmetric dispersal, our model reveals that ecologically-based divergent selection causes nearly complete reproductive isolation between the habitats: successful backcrossing of pure lake deme individuals with admixed individuals occurs only in immediate proximity to the habitat transition, and upstream of that zone, the individuals’ genomic composition rapidly approaches the one of the pure stream deme. The evolution of such strong reproductive isolation in our simulations does not require a phase of physical isolation, but represents primary divergence from standing genetic variation. The simulations further indicate that antagonism between dispersal and selection, and the concomitant oscillations in genomic composition within generations, can persist as an equilibrium, in line with the deep admixture over many generations inferred empirically for most transition zone individuals. In the long run, this equilibrium may be shifted toward stronger reproductive isolation through the accumulation of additional postzygotic barriers. However, given the slow pace of such mutational divergence (e.g., Bolnick & Near 2005; Hedges et al. 2015) and the absence of irreversibly isolated freshwater species within the genus *Gasterosteus*, Misty lake-stream stickleback are likely to become extinct or collapse before complete reproductive isolation has arisen (Taylor et al. 2006; Hendry et al. 2009; Behm et al. 2010; Rosenblum et al. 2012).

### Admixture in the marsh habitat

The main focus of our investigation was the zone over which the main shift in genome-wide allele frequencies occurs – the lowest section of the inlet stream. The temporally replicated (2016, 2017) samples from the site M1 in the adjoining marsh habitat nevertheless allowed a glimpse into the genomic consequences of potential dispersal out of the stream. In both marsh samples, individuals tended toward the genomic composition of the pure lake ecotype. The first sample, however, additionally included individuals showing signs of substantial admixture, a trend much weaker in the sample taken one year later (Fig. 7; a global ancestry analysis including the marsh samples is presented in Fig. S7).

**Fig. 7.**
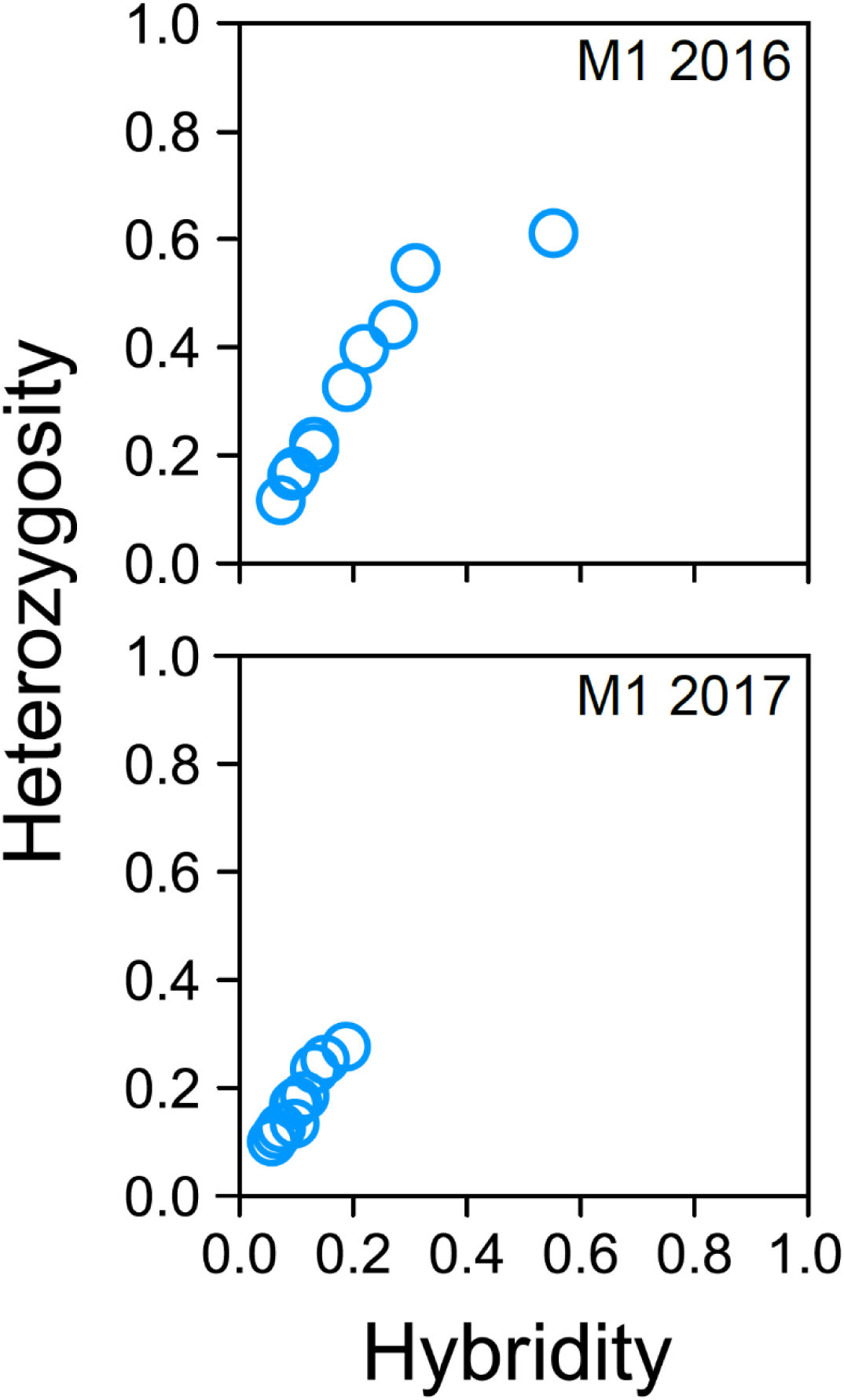
Hybridity and heterozygosity for individuals from the marsh site, sampled in two consecutive years.

Our data from the marsh demonstrate that the lower stream section not only receives immigrant lake ecotypes, but also represents a source of downstream dispersal, as the admixed marsh stickleback must result from interbreeding between the lake ecotype and admixed individuals from the lower stream. The selective conditions in the marsh are not well understood; hence it is unclear to what extent marsh-resident fish would evolve toward the stream ecotype if there were no immigration from the lake. Nevertheless, the genetic composition of marsh stickleback is certainly strongly determined by dispersal from the lake (see also Hanson et al. 2016). The evidence is, first, that pure lake ecotypes still predominate at our lowest stream site S1 located beyond the marsh, hence dispersal from the lake into the marsh should be even more common. Second, the genomic composition of the marsh samples overall resembles the ones obtained in our simulations performed with the highest dispersal rates (Fig. 6B, bottom row right). This particularly holds for the 2017 sample. Indeed, the temporal genomic shift toward the lake ecotype in the marsh can be attributed to strong precipitation after the 2016 sampling period that rose the lake water level, thus flooding the marsh and facilitating dispersal of the lake ecotype (Haenel et al. 2021).

### Conclusions

Lake-stream stickleback pairs represent iconic examples of parapatric divergence and reproductive isolation, yet patterns of hybridization and admixture across their transition zones have not been scrutinized genomically. Here we demonstrate that despite massive dispersal, admixture between the lake and stream ecotypes in the Misty system is largely limited to a narrow zone around the habitat transition. Strong reproductive isolation has here emerged rapidly and without physical barriers as a side-product of adaptive divergence, likely promoted by both ecogeographic and genetic factors: on the one hand, the lake-stream transition is geographically sharp, leading to a steep ecological gradient precluding a niche for phenotypically intermediate hybrid forms (Endler 1977; Nosil et al. 2009). The stream habitat’s linear nature further facilitates adaptive divergence (Gavrilets 2004; Gavrilets & Losos 2009). On the other hand, stickleback fish are known for their abundant standing genetic variation, allowing for highly polygenic adaptive divergence causing reproductive isolation to extend rapidly across the whole genome (Barton 1983; Barton & Bengtsson 1986; Kruuk et al. 1999; Flaxman et al. 2014).

Experimental work on hybrid zones in other organisms is needed address how generally and to what extent divergent ecology is implicated in the buildup and maintenance of steep genomic transitions (Harrison & Larson 2016; Gompert et al. 2017; Moran et al. 2021). It is likely that the rapid emergence of strong reproductive isolation across habitat transitions seen in stickleback fish is unusual, and that steep hybrid zones more typically reflect primarily intrinsic incompatibilities and/or sexual barriers evolved over extended periods of geographic isolation.

## Supporting information

All supplemental information

## Acknowledgements

The study was made possible by financial support from the Swiss National Science Foundation (grant 310030_200374 to DB), and field sampling additionally by Fisheries and Oceans Canada, the British Columbia Ministry of the Environment, and the McGill University Biology Department. Field sampling was aided by Fiona Beaty, Brody Forst, Bailey Feddersen, Tristan Kosciuch, Minxin Lu, Erica MacClaren, Emily McIntosh, Alexanne Oke, Sarah Sanderson, and Mac Willing. Western Forest Products provided logistical and safety support and access to field sites. We thank Ina Nissen and Christian Beisel (Genomics Facility Basel) for advice on sequencing, Russell Corbett-Detig for support in local ancestry estimation, and Lucas Blattner and Lukas Zimmermann for helping run software.

## Author contributions

OB: study design; DNA sample curation; analyses and graphing; interpretation of results; writing of a first manuscript draft. QH: all DNA extractions. KO: field sampling. APH: field sampling; funding. DB: study initiation, design and supervision; funding; analyses, simulations and graphing; interpretation of results; writing of the final manuscript, with feedback from co-authors.

## Data availability

The raw sequence reads are available from the NCBI sequence read archive under the BioProject accession number PRJNA1218656 (https://www.ncbi.nlm.nih.gov/sra/PRJNA1218656). Individual accession numbers are given in Table S1. The full raw genotype matrix from which all files for analysis were derived is provided on the figshare repository (doi: XXXX). A complete compilation of all analytical code is available from the same figshare link.

