## Supplementary material for "Hybridization but minimal introgression: ecologically-based divergent selection maintains a steep hybrid zone in parapatric stickleback fish": All supplemental information

Fig. S1

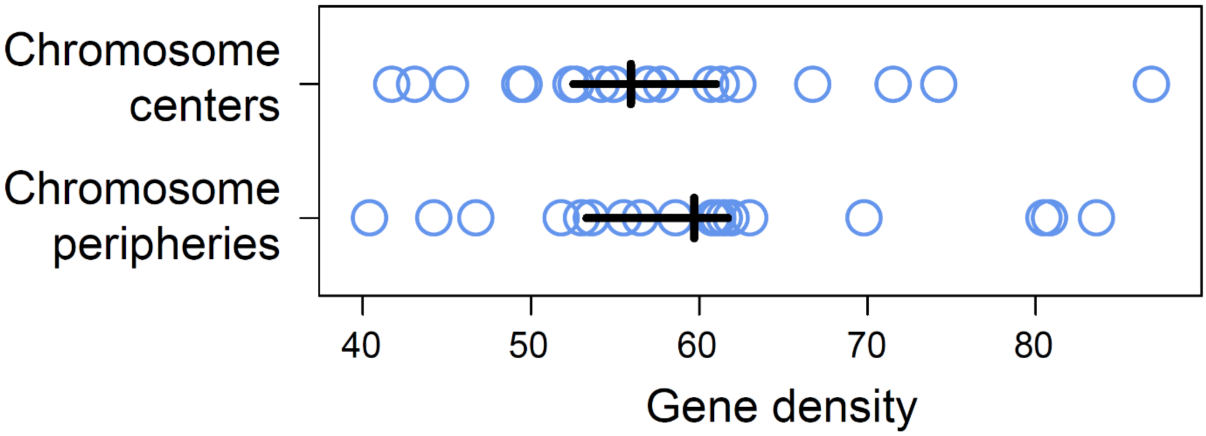


**Fig. S1**. Gene density in the center versus peripheries of the threespine stickleback chromosomes, expressed as the number of genes per megabase. The chromosome peripheries are defined as the terminal 5 Mb on either side of each chromosome. The blue circles represent values for individual chromosomes, the vertical black lines indicate the median across these values, and the horizontal black lines show the 95% compatibility interval for this median based on 10,000 bootstrap resamples. The compatibility intervals are very similar, indicating no substantial difference in gene density between the two chromosome regions.

Fig. S2

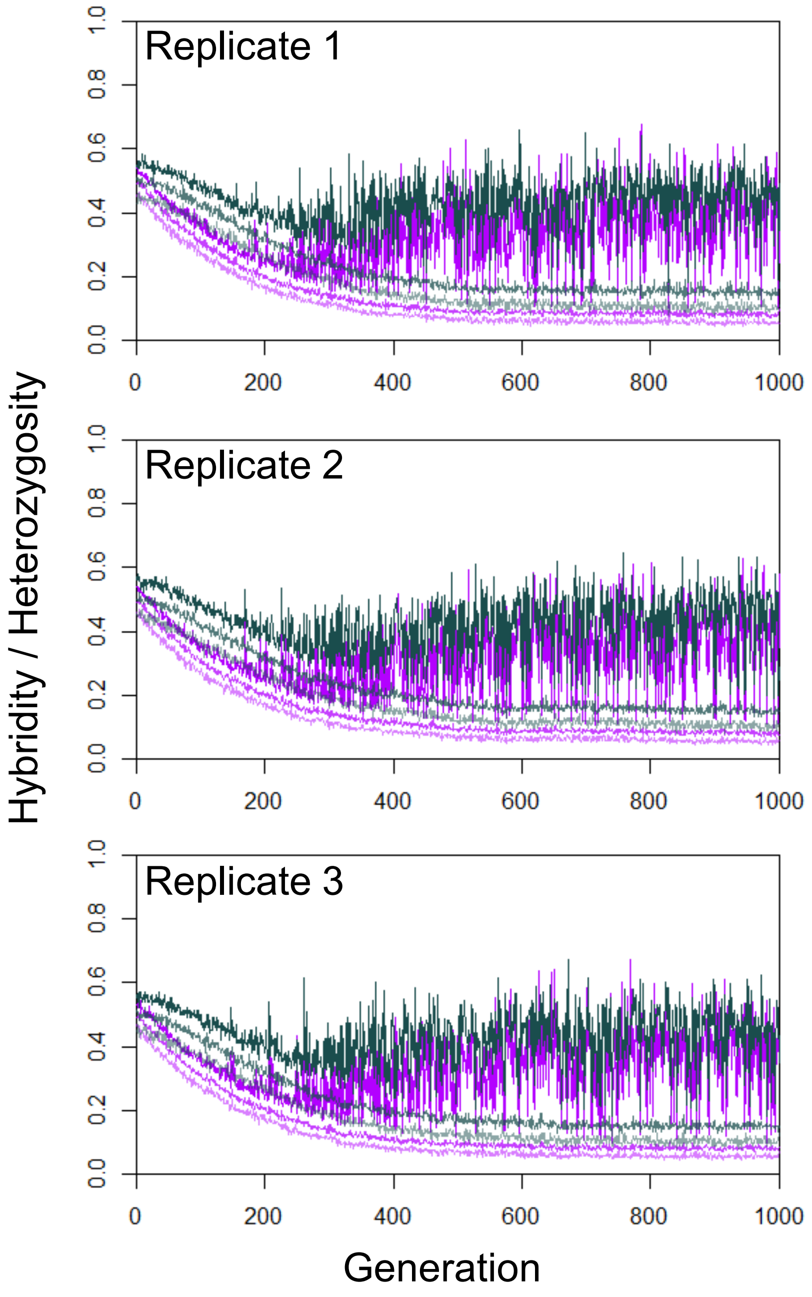


**Fig. S2**. Establishment of migration-selection equilibrium during the simulations of lake-stream divergence with the stepping-stone model using ABC estimation optima for the *f* and *m* parameters. Shown are the 0.125, 0.5, and 0.875 quantiles (separated by increasing color intensities) for hybridity (purple) and heterozygosity (gray) across 40 individuals sampled from the first stream deme in the model in each generation. Results are shown for three replicate simulation runs. Allele are initially sampled at random with a probability of 0.5, hence the genetic composition is initially uniform. After around 500 generations, the system reaches equilibrium, as revealed by the divergence within this deme between lake-adapted immigrants and strongly admixed individuals.

Fig. S3

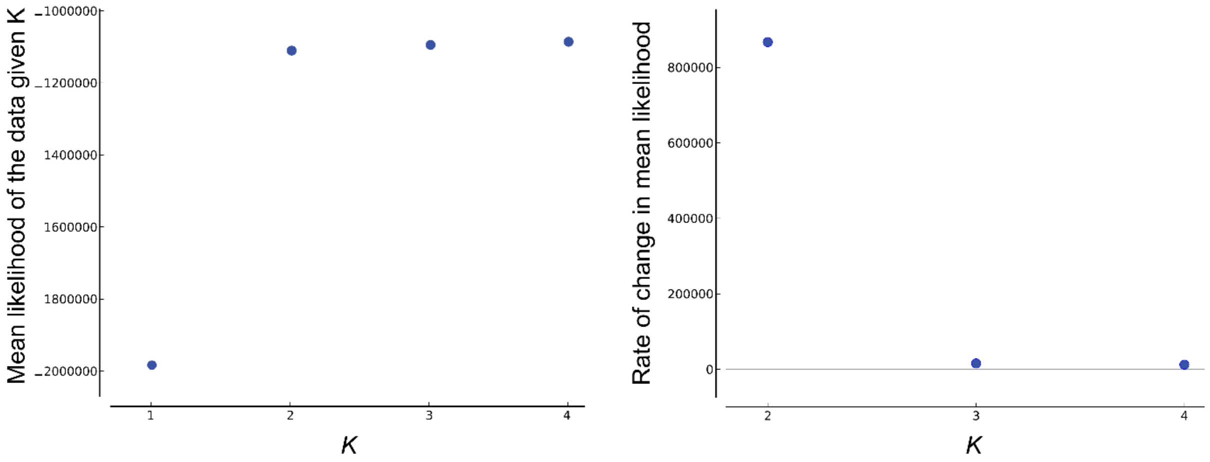


**Fig. S3**. Analysis of population structure with the software STRUCTURE, based on the ecotype-distinctive SNPs and including all 100 individuals from the Misty lake-stream system. The left graphic shows the likelihood *L(K)* of the genomic data when assuming different numbers of populations (*K*), averaged across 20 replicate runs for each *K*. This demonstrates that assuming a single population across the Misty system is implausible, and that beyond *K* = 2, there is minimal gain in likelihood. The rate of change in likelihood *L'(K)* (right graph) thus clearly reveals two true populations (*K* = 2), as does the Δ*K* statistic derived from this metric (not shown).

Fig. S4

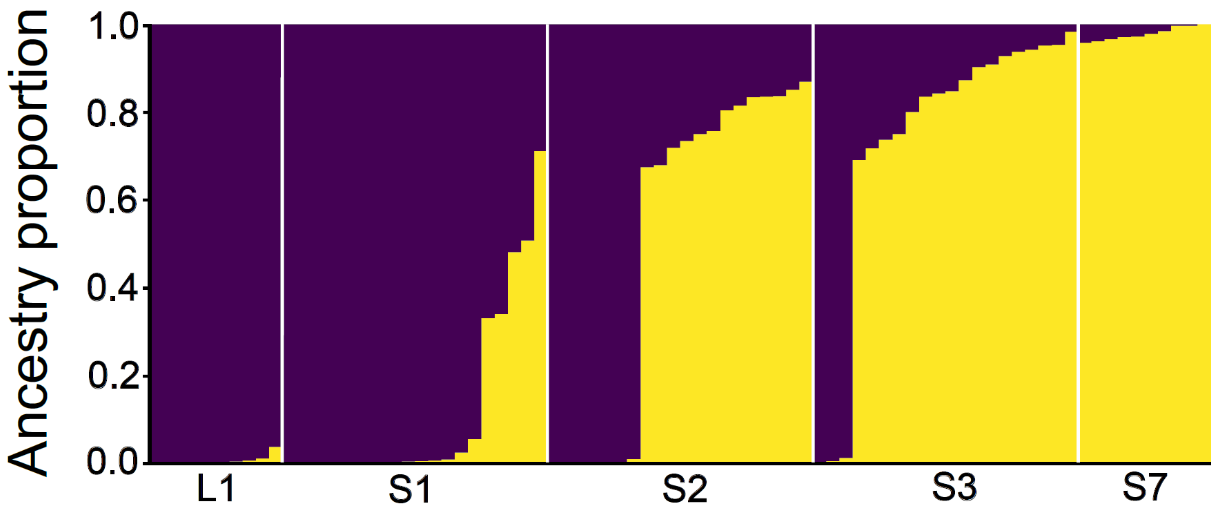


**Fig. S4**. Global ancestry proportions as estimated by STRUCTURE. The graph follows the conventions of Fig. 2B, except that the underlying markers are random SNPs, not ecotype-distinctive ones.

Fig. S5

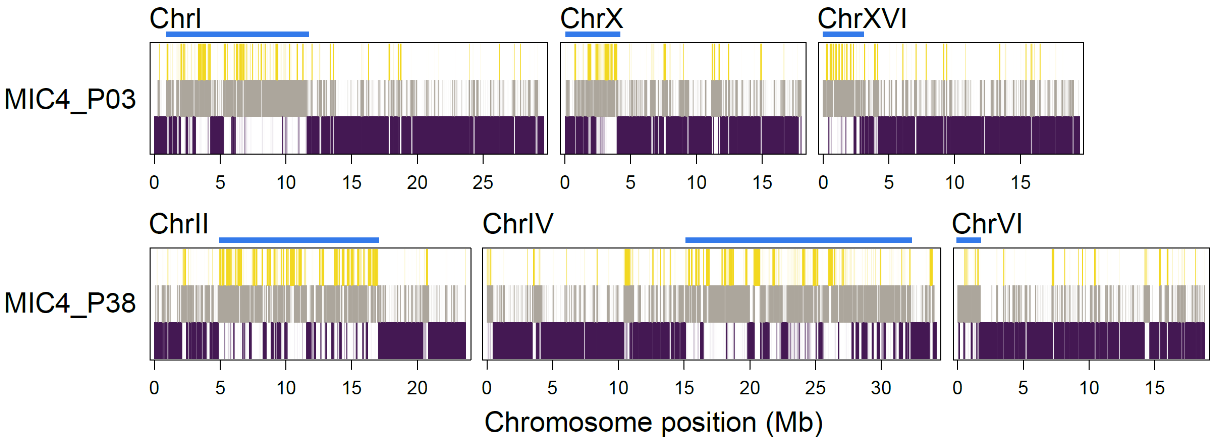


**Fig. S5**. Two individuals from the first stream site (S1) showing tracts of alternative ancestry along some chromosomes. Specifically, while most chromosomes in these individuals exhibited predominantly homozygous lake ancestry (like in Fig. 4 top row), a few chromosomes displayed extended tracts of heterozygous ancestry (approximately indicated by horizontal blue bars), suggesting relatively recent backcrossing of dispersers from the lake with admixed local individuals.

Fig. S6

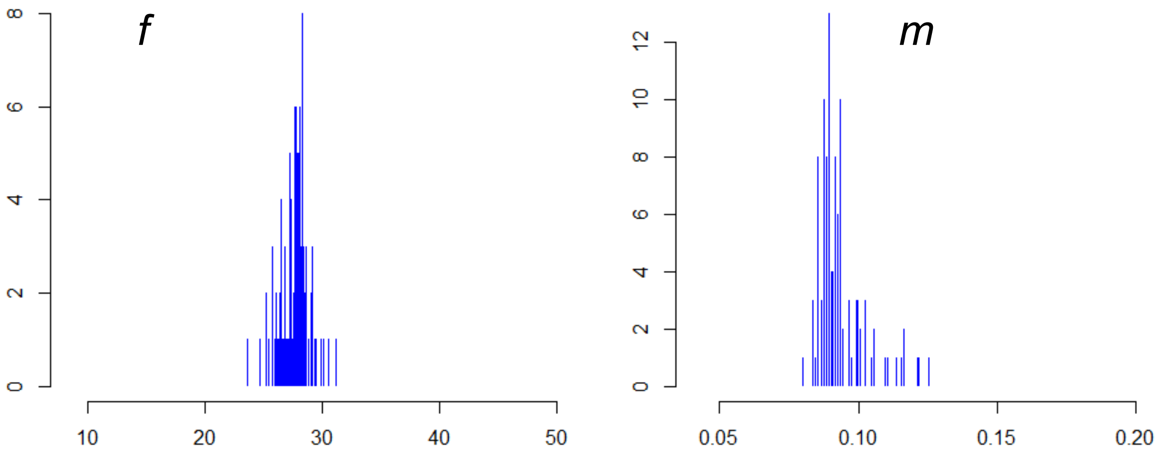


**Fig. S6**. Distribution of Approximate Bayesian Computation (ABC) parameter estimates for the factor *f* determining the size of the lake deme relative to the size of the stream demes (*n* = 200), and for the proportion *m* of individuals dispersing in each generation from a given deme into each neighboring deme (except for the terminal demes, total emigration is thus 2*m*). The distributions are based on 100 replicate ABC estimation runs, all performed with hybridity and heterozygosity summary statistics from 300 simulations with values of *f* and *m* chosen at random. Shown are median estimates from each estimation run. The grand medians (27.8 and 0.091 for *f* and *m*) were taken as optima for further simulation.

Fig. S7

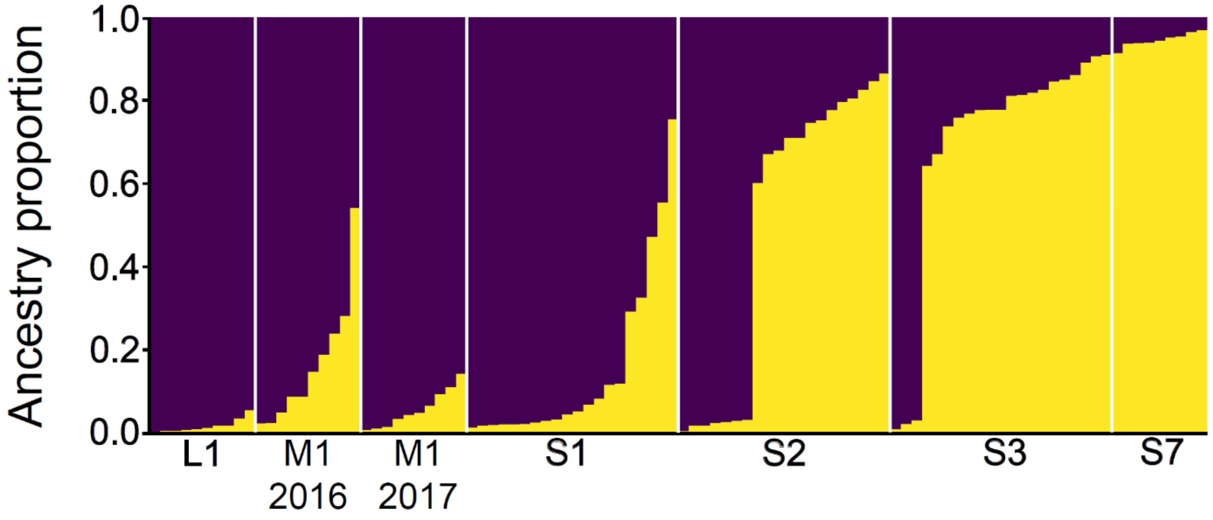


**Fig. S7**. Global ancestry proportions as estimated by STRUCTURE based on the ecotype-distinctive markers. The graph follows the conventions of Fig. 2B, but includes the individuals from the two temporal marsh site (M1) samples (*n* = 10 each).
